# Evidence for vagal sensory neural involvement in influenza pathogenesis and disease

**DOI:** 10.1101/2023.08.29.555274

**Authors:** Nathalie A.J. Verzele, Brendon Y. Chua, Kirsty R. Short, Aung Aung Kywe Moe, Isaac N. Edwards, Helle Bielefeldt-Ohmann, Katina D. Hulme, Ellesandra C. Noye, Marcus Z.W. Tong, Patrick C. Reading, Matthew W. Trewella, Stuart B. Mazzone, Alice E. McGovern

**Affiliations:** School of Chemistry and Molecular Biosciences, The University of Queensland, St Lucia Queensland 4072, Australia; Department of Anatomy and Physiology, The University of Melbourne, Parkville, Victoria 3010, Australia; The Peter Doherty Institute for Infection and Immunity, Department of Microbiology and Immunology, University of Melbourne, Melbourne, Victoria 3000, Australia; Australian Infectious Diseases Research Centre, The University of Queensland, St Lucia, Queensland 4072, Australia; Department of Medical Imaging and Radiation Sciences, Monash University, Clayton, Victoria 3800, Australia; WHO Collaborating Centre for Reference and Research on Influenza, Victorian Infectious Disease Reference Laboratory, Peter Doherty Institute for Infection, and Immunity, 792 Elizabeth St., Melbourne, VIC 3000, Australia

## Abstract

Influenza A virus (IAV) is a common respiratory pathogen and a global cause of significant and often severe morbidity. Although inflammatory immune responses to IAV infections are well described, little is known about how neuroimmune processes contribute to IAV pathogenesis. In the present study, we employed surgical, genetic, and pharmacological approaches to manipulate pulmonary vagal sensory neuron innervation and activity in the lungs to explore potential crosstalk between pulmonary sensory neurons and immune processes. Intranasal inoculation of mice with H1N1 strains of IAV resulted in stereotypical antiviral lung inflammation and tissue pathology, changes in breathing, loss of body weight and other clinical signs of severe IAV disease. Unilateral cervical vagotomy and genetic ablation of pulmonary vagal sensory neurons had a moderate effect on the pulmonary inflammation induced by IAV infection, but significantly worsened clinical disease presentation. Inhibition of pulmonary vagal sensory neuron activity via inhalation of the charged sodium channel blocker, QX-314, resulted in a moderate decrease in lung pathology, but again this was accompanied by a paradoxical worsening of clinical signs. Notably, vagal sensory ganglia neuroinflammation was induced by IAV infection and this was significantly potentiated by QX-314 administration. This vagal ganglia hyperinflammation was characterized by alterations in IAV-induced host defense gene expression, increased neuropeptide gene and protein expression, and an increase in the number of inflammatory cells present within the ganglia. These data suggest that pulmonary vagal sensory neurons play a role in the regulation of the inflammatory process during IAV infection and suggest that vagal neuroinflammation may be an important contributor to IAV pathogenesis and clinical presentation. Targeting these pathways could offer therapeutic opportunities to treat IAV-induced morbidity and mortality.

**Author summary:** Influenza viruses are a common respiratory pathogen that represent a constant and pervasive threat to human health. Although the inflammatory and immune responses to influenza viral infections are well described, little is known about the role the nervous system plays in the formation and progression of disease. The lungs receive a rich supply of sensory nerve fibers from the vagus nerve. These nerves are critical for protecting the lungs against harmful stimuli and play an important defence role against pathogens, including viruses. Here we use several complex animal models to demonstrate the impact lung sensory neurons have on influenza viral infection and disease outcome. We demonstrate that ablation of lung sensory neurons and inhibition of their neural activity significantly worsens the clinical outcome in mice infected with influenza virus, however with only a moderate impact on lung pathology. Interestingly, when the activity of these neurons is inhibited during influenza viral infection, this drives a hyper neuroinflammatory response within the vagal sensory ganglia, where their cell bodies are located. Our work provides new insights into how these lung sensory neurons are involved in influenza viral infections and may offer therapeutic opportunities to treat influenza-induced morbidity and mortality.

## Introduction

Influenza A virus (IAV) is a common and highly contagious respiratory virus, causing significant morbidity and mortality in humans worldwide [1, 2]. Mild IAV infections present with a range of symptoms including a sudden onset of fever, chills, sneeze, cough, congestion, headaches, malaise, and lethargy which generally resolve soon after viral clearance [1, 2]. However, in some individuals, including those considered at risk, infections can be life threatening resulting in clinically severe IAV disease associated with pulmonary edema, hypoxemia, pneumonia, and acute respiratory distress syndrome with symptomatology persisting well beyond the clearance of the viral infection.

Although it is well accepted that the pulmonary inflammatory response to IAV is mediated by a diverse range of both resident and recruited cells, much of this *a priori* knowledge fails to recognize the important role played by the nervous system, and in particular sensory neurons, in pulmonary inflammation and disease symptomology. From the larynx to the lung parenchyma, the mammalian respiratory system is innervated by sensory neurons that are almost exclusively derived from the vagus nerve. These vagal sensory neurons are critical for monitoring the pulmonary environment and upon activation transmit information to the brainstem that is then used to drive reflexes and respiratory behaviours [3, 4]. These sensory neurons are not homogeneous in phenotype, but rather different subtypes exist that are classified based on their neurochemistry, molecular expression profiles, physiological properties, and the reflexes that they initiate [4-7].

Some vagal sensory neurons are responsive to potentially damaging stimuli including inhaled, aspirated or locally produced chemicals. These sensory neurons often express receptors for a wide range of immune cell derived molecules [4-6, 8] and can be directly activated or modulated by inflammation [4, 7-12]. In this regard, vagal sensory neural pathways may be particularly important in the generation of symptoms during respiratory viral infections as they provide input to diverse brain neural circuits that regulate respiratory reflexes, breathing, mood and other complex behaviors [13-18]. We have previously shown that during an acute infection with IAV, the pulmonary vagal sensory neurons undergo transcriptional changes and take on a neuroinflammatory phenotype. This neuropathy is characterized by an infiltration of immune cells into the vagus nerve and the upregulation of genes in the sensory neurons associated with host defense and inflammation [19]. In addition, these neurons respond by translocating the alarmin high mobility group box-1 (HMGB1) from the nucleus to the cytoplasm, which could serve as a mediator of hyperinnervation and prolonged hypersensitivity by promoting vagal sensory neurite growth and excitability [20].

Interactions between pathogens, inflammation and sensory nerves are unlikely to be unidirectional and recent studies have shown that the nervous system can reciprocally regulate immune processes associated with respiratory disease, notably via pathways involving neuropeptide-containing pulmonary sensory neurons [21-23]. Here, we investigated this interaction further using a combination of surgical, genetic, and pharmacological approaches to assess the effects of interrupting pulmonary sensory neuroimmune interactions on IAV pathogenesis, vagal neuroinflammation and disease.

## Results

### Vagal denervation impacts IAV pathogenesis and disease

We initially sought to characterize the relative contribution of left and right vagus nerves with respect to the sensory innervation to left and right lungs (Fig 1A). Following injection of the retrograde serotype adeno-associated virus encoding tdTomato (AAVrg^TdT^) into the left lung, 75.4% of labelled neurons were in the left vagal ganglia while 24.6% in the right vagal ganglia (Fig 1B and B’), indicating the left lung receives significantly more sensory innervation from the left compared to the right vagal ganglia. In comparison, the vagal sensory neurons contributing to the right lung appeared to be more evenly distributed across right (59.1%) and left (40.9%) vagal ganglia. Unilateral cervical vagotomies eliminated AAVrg^TdT^ traced neurons in the ipsilateral ganglia only. In all experiments included in the analysis, the injection site was contained to the injected lung, with no observable spread of AAVrg^TdT^ into the contralateral lung or pleural cavity (data not shown).

**Figure 1.**
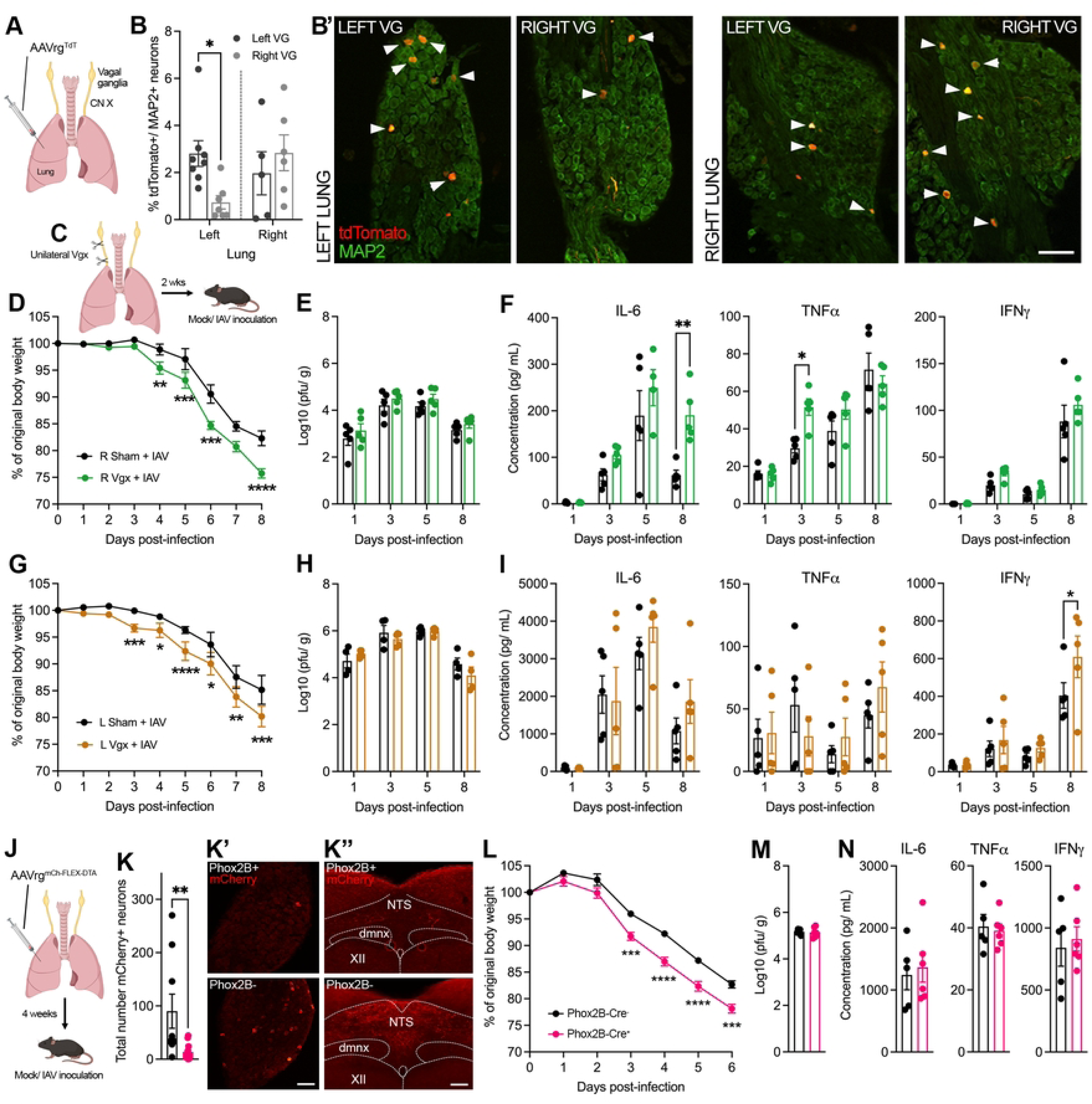
Vagal denervation impacts IAV pathogenesis and disease related to IAV disease. (A) Schematic showing retrograde viral tracing to quantify the ratio of right and left lung projecting sensory neurons in the ipsilateral versus contralateral vagal sensory ganglia. (B) Quantification of the percentage of tdTomato+ neurons per MAP2+ neurons in either left or right vagal sensory ganglia innervating the left (n = 8) or right lung (n = 6) with (B’) representative images depicting examples of traced neurons in the vagal sensory ganglia. (C) Schematic outlining the procedure for surgically removing partial vagal innervation to the lungs (right or left unilateral vagotomy). Graphs depict group level body weight change, lung viral titers and lung cytokine measurements (n = 5 per group at days 1, 3, 5, 8 post IAV or mock infection) over the course of IAV infection following either (D, E, F) right or (G, H, I) left vagotomy or sham surgery. (J) Schematic showing genetic ablation of nodose lung sensory neurons in the vagal sensory ganglia using retrograde AAV encoding Cre-inducible diphtheria toxin (AAVrg mCherry-FLEX-DTA) in Phox2B-Cre^+^ (n = 6) and Phox2B-Cre**^-^** (n = 5) mice. (K) Quantification of the total number of mCherry+ neurons in the vagal sensory ganglia with (K’ and K”) showing representative images of the vagal sensory ganglia and brainstem nucleus of the solitary tract (NTS) taken from Phox2B-Cre^+^ (top panel) and Phox2B-Cre^-^ (bottom panel) mice. Graphs depict group level (L) body weight change, (M) lung viral titers and (N) lung cytokine measurements over the course of IAV infection following nodose lung sensory neuron ablation. Data represented as mean ± SEM. *, **, ***, **** denotes significance of *p* < 0.05, *p* < 0.01, *p* < 0.001, *p* < 0.0001, respectively, as (B) determined by two-way ANOVA corrected for multiple comparisons (Šídák), (D, F, G, I, L) repeated measures two-way ANOVA corrected for multiple comparisons (Šídák) and (K) Mann-Whitney t-test. CN X, vagus nerve; VG, vagal ganglia; Vgx, vagotomy; dmnx, dorsal motor nucleus of the vagus; XII, hypoglossal motor nucleus. Scale bar represents 100μm.

Considering these findings, we opted to perform both right and left unilateral vagotomies in separate animal cohorts and compared the results to sham vagotomized controls (Fig 1C). Surgical removal of a portion of either the right or left cervical vagus resulted in an initial and transient reduction in body weight compared to sham controls (data not shown). Once mice returned to their pre-surgical body weight (∼2 weeks) they were intranasally inoculated with 50 plaque forming units (PFU) of the mouse adapted H1N1 IAV, strain A/Puerto Rico/8/34 (PR8). Control (mock infected) mice received intranasal phosphate buffered saline (PBS). No differences in bodyweight were observed in mock infected mice of both right/ left sham and right/ left unilateral vagotomized groups (Supp Fig 1A and B). IAV infected mice (both right/ left sham and vagotomized groups) began to lose body weight around days 3 to 4 post-infection, with vagotomized mice losing significantly more weight than their sham IAV infected counterparts (Fig 1D and G).

Lung viral titers remained consistent between both right/ left vagotomized and sham vagotomized IAV infected mice (Fig 1E and H). We did not observe any differences between lung cytokines and inflammatory cell infiltrate between mock infected mice of both right/ left sham and right/ left vagotomized groups (Supp Fig 1C and D; Tables 1 and 2). The induction of cytokines (IL-6, IFNγ, TNFα) observed in lung lysates of right/ left vagotomized IAV infected mice showed some differences compared to sham vagotomized IAV infected mice. TNFα and IL-6 were significantly lower in right vagotomized mice at days 3 and 8 post-infection, respectively (TNFα: vagotomized 51.6 ± 4.4 pg/ ml, sham 29.6 ± 2.4 pg/ ml, day 3, *p* = 0.0107; IL-6: vagotomized 190.6 ± 26.0 pg/ ml, sham 61.6 ± 10.7 pg/ ml, day 8, *p* = 0.0036; Fig 1F), and left vagotomized mice had an increase in IFNγ at day 8 post-infection (vagotomized: 609.6 ± 110.6 pg /mL, sham: 404.2 ± 68.1 pg/mL, *p* = 0.028; Fig 1I). In addition, we observed occasional differences between immune cells in the bronchoalveolar lavage fluid (BALF) of both right/ left vagotomized IAV infected mice compared to their sham groups. Of note, natural killer (NK) T-cells were more frequent in both right and left vagotomized IAV infected mice at day 8 post-infection (*p* = 0.0414 and *p* = 0.0496, respectively; Tables 1 and 2). CD4^+^ and CD8^+^ T-cells were significantly decreased in right vagotomized IAV infected mice at day 8 post-infection compared to their sham group (*p* = 0.0032 and *p* = 0.0139, respectively; Table 1), whereas for left vagotomized IAV infected mice we found an increase in CD8^+^ T-cells (*p* = 0.0021; Table 2). No difference was observed in systemic cytokine levels between IAV infected vagotomized and sham mice (Table 3).

**Table 1.** Flow cytometry of BALF from unilateral right vagotomized and sham groups post-infection.

|  | Mock |  | Day 1 PI |  | Day 3 PI |  | Day 5 PI |  | Day 8 PI |  |
| --- | --- | --- | --- | --- | --- | --- | --- | --- | --- | --- |
|  | Sham | Vgx | Sham | Vgx | Sham | Vgx | Sham | Vgx | Sham | Vgx |
| Neutrophils | 0.83 ± 0.17 | 0.54 ± 0.13 | 4.2 ± 1.0 | 3.4 ± 0.5 | 13.5 ± 3.0 | 21.6 ± 2.9 | 47.2 ± 6.9 | 72.8 ± 11.7 | 5.0 ± 0.4 | 9.8 ± 1.2 |
| Dendritic cells | 40.7 ± 7.3 | 40.8 ± 2.1 | 52.9 ± 9.2 | 66.7 ± 8.6 | 54.9 ± 2.0 | 53.2 ± 2.9 | 165.9 ± 15.3 | 188.7 ± 28.8 | 441.9 ± 61.3 | 366.1 ± 32.2 |
| Alveolar macrophages | 0.72 ± 0.17 | 0.63 ± 0.14 | 2.8 ± 0.6 | 2.4 ± 0.4 | 0.70 ± 0.07 | 0.93 ± 0.14 | 19.0 ± 1.5 | 13.2 ± 3.0 | 8.3 ± 1.3 | 4.3 ± 0.3 |
| Interstitial macrophages | 33.6 ± 6.0 | 38.2 ± 4.4 | 55.3 ± 10.8 | 68.9 ± 11.4 | 79.3 ± 12.0 | 64.6 ± 7.4 | 172.6 ± 18.5 | 158.6 ± 17.8 | 121.6 ± 16.0 | 225.4 ± 36.4 |
| B-cells | 284.5 ± 49.4 | 334.5 ± 19.3 | 331.5 ± 55.1 | 419.9 ± 64.7 | 317.3 ± 10.4 | 220.0 ± 22.3* | 376.8 ± 16.1 | 315.5 ± 39.4 | 728.2 ± 135.7 | 588.7 ± 43.7 |
| NK-cells | 140.9 ± 24.6 | 167.2 ± 8.0 | 139.2 ± 22.4 | 178.1 ± 25.3 | 189.6 ± 11.2 | 166.4 ± 13.6 | 406.0 ± 32.2 | 360.0 ± 40.9 | 481.5 ± 52.8 | 619.5 ± 61.1 |
| CD4<sup>+</sup> T-cells | 78.3 ± 11.6 | 85.0 ± 5.5 | 86.4 ± 14.8 | 106.0 ± 16.2 | 92.4 ± 4.3 | 76.0 ± 8.2 | 116.4 ± 5.7 | 91.5 ± 10.5 | 833.4 ± 130.7 | 607.6 ± 25.7** |
| NKT-cells | 20.7 ± 8.4 | 20.1 ± 8.2 | 45.9 ± 18.8 | 58.7 ± 23.9 | 183.3 ± 74.8 | 180.0 ± 73.5 | 196.1 ± 80.1 | 237.4 ± 96.9 | 282.5 ± 115.3 | 627.9 ± 256.4*** |
| CD8<sup>+</sup> T-cells | 55.9 ± 9.0 | 70.1 ± 4.7 | 59.6 ± 9.9 | 70.3 ± 11.6 | 71.8 ± 2.5 | 56.0 ± 6.6 | 73.4 ± 5.6 | 66.8 ± 10.3 | 1205.8 ± 159.4 | 951.4 ± 77.0* |

Values are x10<sup>3</sup> total number of cells. Data represented as the mean ± SEM. \**p* = < 0.05, \*\**p* = < 0.01, \*\*\**p* = < 0.001, Vagotomy (Vgx) versus
Sham as determined by repeated measures 2-way ANOVA Tukey's multiple comparisons test. N = 5/ group. PI, post-infection; NK, natural killer.

**Table 2.** Flow cytometry of BALF from unilateral left vagotomized and sham groups post-infection.

|  | Mock |  | Day 1 PI |  | Day 3 PI |  | Day 5 PI |  | Day 8 PI |  |
| --- | --- | --- | --- | --- | --- | --- | --- | --- | --- | --- |
|  | Sham | Vgx | Sham | Vgx | Sham | Vgx | Sham | Vgx | Sham | Vgx |
| Neutrophils | 0 ± 0 | 3 ± 3 | 4 ± 1 | 6 ± 3 | 691 ± 423 | 837 ± 694 | 330 ± 150 | 712 ± 453 | 371 ± 188 | 761 ± 160 |
| Dendritic cells | 2 ± 0 | 4 ± 0 | 0 ± 0 | 0 ± 0 | 15 ± 7 | 16 ± 6 | 35 ± 11 | 10 ± 4 | 92 ± 28 | 76 ± 29 |
| Alveolar macrophages | 1761 ± 518 | 1050 ± 165 | 788 ± 219 | 873 ± 181 | 706 ± 257 | 1043 ± 298 | 644 ± 154 | 845 ± 98 | 492 ± 46 | 736 ± 157 |
| Interstitial macrophages | 1 ± 1 | 2 ± 1 | 11 ± 7 | 8 ± 3 | 1043 ± 661 | 647 ± 463 | 357 ± 91 | 903 ± 508 | 148 ± 67 | 401 ± 98 |
| B-cells | 3 ± 1 | 2 ± 1 | 3 ± 2 | 7 ± 3 | 111 ± 33 | 147 ± 122 | 115 ± 40 | 248 ± 127 | 193 ± 69 | 494 ± 72* |
| NK-cells | 1489 ± 526 | 1386 ± 308 | 462 ± 156 | 691 ± 215 | 2259 ± 750 | 1979 ± 1164 | 782 ± 240 | 1860 ± 827 | 998 ± 228 | 3154 ± 820 |
| CD4<sup>+</sup> T-cells | 4 ± 2 | 5 ± 1 | 0 ± 0 | 1 ± 0 | 28 ± 11 | 20 ± 10 | 111 ± 46 | 95 ± 46 | 1274 ± 211 | 1401 ± 490 |
| NKT-cells | 0 ± 0 | 0 ± 0 | 0 ± 0 | 0 ± 0 | 50 ± 36 | 16 ± 11 | 161 ± 33 | 74 ± 32 | 150 ± 31 | 353 ± 169* |
| CD8<sup>+</sup> T-cells | 7 ± 3 | 7 ± 2 | 0 ± 0 | 0 ± 0 | 40 ± 23 | 27 ± 14 | 334 ± 150 | 135 ± 91 | 2777 ± 588 | 6342 ± 1474** |

Data represented as the mean ± SEM. \* $p < 0.05$ , \*\* $p < 0.01$ , Vagotomy (Vgx) versus Sham as determined by repeated measures 2-way ANOVA
multiple comparisons test. N = 5/ group. PI, post-infection; NK, natural killer

**Table 3.** Serum cytokines following vagotomy and IAV infection.

|  | Mock |  | Day 1 PI |  | Day 3 PI |  | Day 5 PI |  |
| --- | --- | --- | --- | --- | --- | --- | --- | --- |
|  | Sham | Vgx | Sham | Vgx | Sham | Vgx | Sham | Vgx |
| IL-6 | 211.2 ± 42.8 | 289.9 ± 39.1 | 165.8 ± 13.8 | 226.4 ± 13.0 | 402.2 ± 52.8 | 591.9 ± 115.6 | 328.9 ± 48.8 | 425.1 ± 62.8 |
| IL-12 | 304.3 ± 12.1 | 163 ± 6 8.4 | 168.7 ± 71.6 | 263.2 ± 42.4 | 429.5 ± 90.4 | 279.6 ± 57.0 | 497.7 ± 62.2 | 432.6 ± 32.3 |
| TNFα | 55.9 ± 4.1 | 59.6 ± 2.9 | 59.6 ± 2.1 | 60.8 ± 5.4 | 63.9 ± 4.3 | 83.2 ± 10.7 | 65.1 ± 3.7 | 61.1 ± 3.1 |

Data presented as pg/ ml and mean ± SEM. N = 5/ group.

Although the majority of axons comprising the vagus nerve are derived from sensory neurons, a sizeable portion (up to 40%) of the vagal lung innervation originates from parasympathetic motor neurons [24]. In addition, the vagus is not solely a pulmonary nerve but rather innervates numerous other viscera. Therefore, vagotomy is a non-specific method of targeting the sensory innervation supplying the lungs. For more specificity, we developed a genetic approach to selectively ablate lung projecting vagal sensory neurons. In rodents, the majority of sensory innervation in the lungs arise from the nodose portion of the vagal sensory ganglia [4, 25, 26]. The homeodomain transcription factor Phox2B is expressed in nodose neurons and is not present in other neurons that make up the vagal sensory ganglia [27, 28]. To ablate the nodose neurons innervating the lungs, we injected a retrograde AAV encoding Cre-recombinase (Cre) dependent diphtheria toxin A (DTA) (AAVrg^mCh-FLEX-DTA^) into the lungs of Phox2B Cre-expressing (Phox2B-Cre^+^) mice (Fig 1J). AAVrg^mCh-FLEX-DTA^ expresses DTA in Cre-expressing neurons and mCherry in neurons that do not express Cre. Following 4 weeks recovery, we observed a significantly lower number of mCherry expressing neurons in Phox2B-Cre^+^ mice compared to the Phox2B-Cre**^-^** littermates (13.2 ± 4.0 and 90.1 ± 32.0 mCherry neurons, respectively; *p* = 0.0058; Fig 1K and K’). We also observed less mCherry-expressing nerve fibers in the nucleus of the solitary tract (NTS), the brainstem location where nodose afferents centrally project [15, 16, 29] in Phox2B-Cre^+^ mice compared to Phox2B-Cre^-^ mice (Fig 1K”) indicative of correct AAVrg^mCh-FLEX-DTA^ functionality. Similar to surgical vagotomy, Phox2B-Cre^+^ mice receiving intrapulmonary injection of AAVrg^mCh-FLEX-DTA^ lost significantly more body weight compared to the Phox2B-Cre^-^ AAVrg^mCh-FLEX-DTA^ group following IAV infection (Fig 1L). Again, this loss in body weight was not associated with differences observed in lung viral titers (Fig 1M), inflammatory cytokines (Fig 1N) or immune cell infiltrates (Table 4).

**Table 4.** Flow cytometry of BALF from DTA ablated Phox2B lung vagal ganglia neurons of Phox2B-Cre^-^ (N = 5) and Phox2B-Cre^+^ (N = 6) mice at day 8 post-infection.

|  | Phox2B-Cre<sup>-</sup> | Phox2B-Cre<sup>+</sup> |
| --- | --- | --- |
| Neutrophils | 1982.0 ± 283.0 | 1500.0 ± 169.8 |
| Dendritic cells | 448.6 ± 105.0 | 339.0 ± 51.8 |
| Alveolar macrophages | 138.2 ± 33.5 | 125.9 ± 39.8 |
| Interstitial macrophages | 94.9 ± 3.0 | 91.6 ± 82.8 |
| B-cells | 369.2 ± 63.0 | 452.2 ± 79.2 |
| NK-cells | 3028.0 ± 320.5 | 2712.0 ± 196.4 |
| CD4<sup>+</sup> T-cells | 343.2 ± 64.5 | 353.7 ± 58.2 |
| NKT-cells | 32.5 ± 6.6 | 45.4 ± 15.2 |
| CD8<sup>+</sup> T-cells | 1003.0 ± 239.3 | 952.8 ± 158.7 |

Data represented as the mean ± SEM

The results of the unilateral vagotomy and AAVrg^mCh-FLEX-DTA^ experiments suggest that disruption of the vagal sensory innervation to the lungs impacts the clinical presentation of IAV disease severity. Although modest effects on IAV-induced pulmonary pathology are evident when vagal sensory innervation to the lungs is reduced, it is questionable if these changes in pulmonary pathology are the principal cause of the altered clinical presentation.

### Pharmacological inhibition of the activity of pulmonary sensory neurons during IAV infection

Since surgically denervating and ablating sensory neurons prior to IAV infection may conceivably cause changes in the respiratory, immune, and nervous systems, we shifted to using a pharmacological approach to temporarily inhibit lung afferent activity during IAV infection. We also expanded the range of endpoints measured. QX-314 (N-ethyl-lidocaine) is a charged sodium channel inhibitor that enters certain neurons through large-pore ion channels, notably transient receptor potential vanilloid and ankyrin 1 (TRPV1 and TRPA1). These channels are selectively expressed in populations of vagal sensory neurons involved in airway defense and effectively gated (opened) by products of inflammation. Consequently, QX-314 has been used to inhibit the peripheral induction of action potential formation in sensory neurons in a range of pulmonary inflammatory conditions [21, 30, 31]. Mice were inoculated with either mock or the H1N1 IAV strain (Auck/09, 5.5 x 10^3^ PFU) and nebulized with 300μM of QX-314 or vehicle (sterile saline), twice-daily from 3 days post-infection, during the early inflammatory phase when sensory nerve fiber TRPV1/ A1 channels are likely open (Fig 2A) [21, 32, 33]. Importantly, we observed no difference in body weight and clinical score (Supp Fig 2A and B) in mock infected groups receiving either QX-314 or vehicle (PBS). Additionally, baseline respiratory function remained unchanged over the course of the experiment between both mock infected groups (Table 5A-C). In contrast, IAV infected mice exhibited a decrease in body weight and increase in clinical signs, with the IAV infected QX-314 group displaying more significant changes compared to the IAV infected vehicle group (Fig 2B and C). IAV infection caused significant changes in respiratory function in both vehicle and QX-314 mice compared to mock infected mice. As disease progressed, we observed decreases in respiratory rate and relaxation time, increases in tidal volume, peak inspiratory and expiratory flow (PIF and PEF), ejection fraction 50 (EF50), pause and enhanced pause (PAU, Penh) (Tables 5A-C). In IAV infected QX-314 mice, we observed significant increases in EF50, pause and Penh and a trend in increased PEF (Fig 2D; Tables 5A-C) compared to IAV infected vehicle mice. EF50 is the flow rate at which 50% of tidal volume has been expelled and has been shown to increase along with disease severity [34]. Penh is a non-specific measurement of breathing and is calculated based on = (PEF/ PIF) x pause. Therefore, the increases observed in PEF, pause and PIF remaining unchanged likely lead to the increases seen in Penh. Again, this measurement has been shown to increase along with disease severity [34].

**Figure 2.**
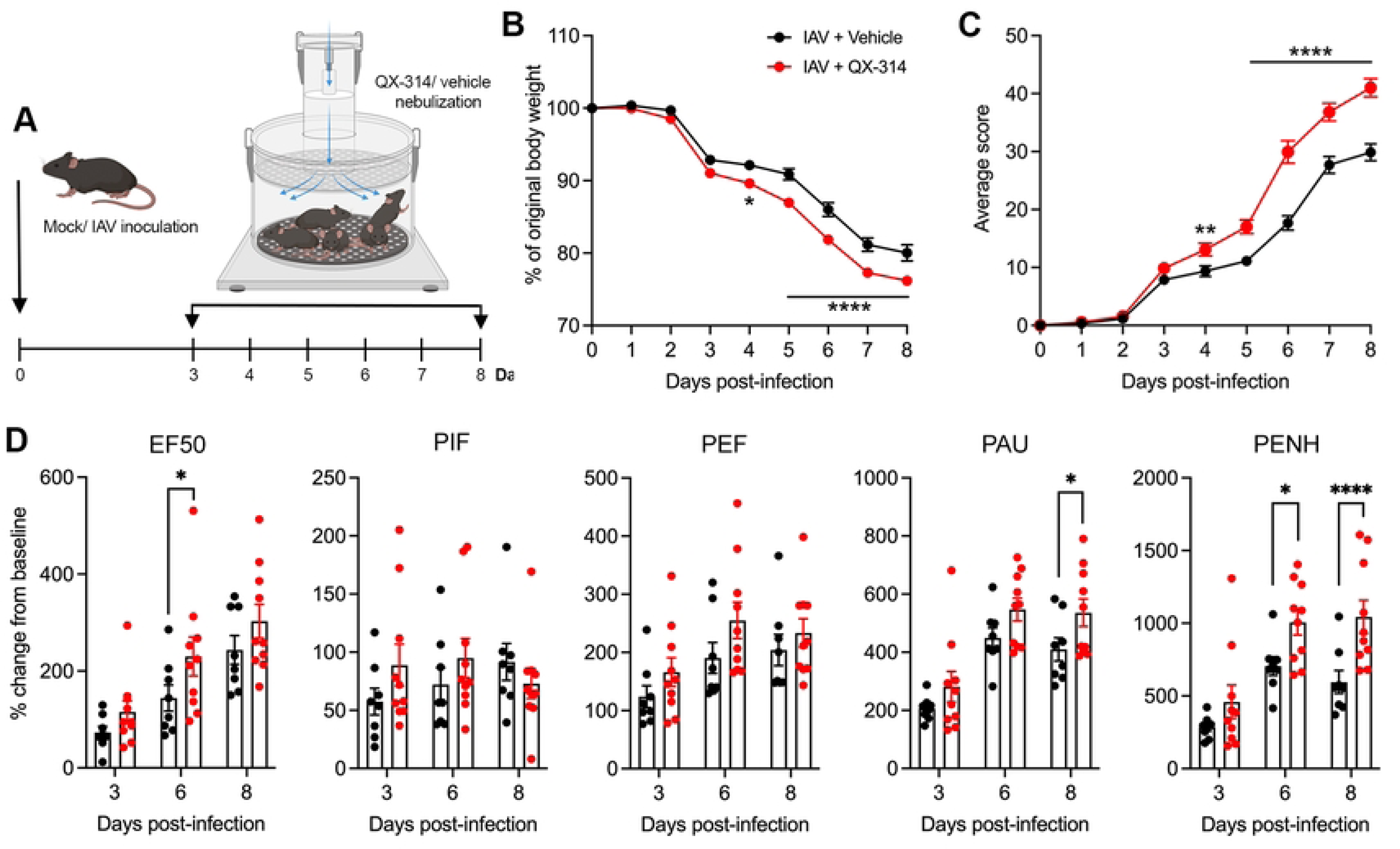
Pharmacological inhibition of pulmonary sensory neuron activity during IAV infection increases morbidity. (A) Schematic outlining experimental design used to assess the effect of nebulized QX-314 on IAV morbidity. Graphs depict group level (B) body weight change, (C) clinical scoring and (D) respiratory parameters of interest post infection with IAV following administration of QX-314 or vehicle. N = 10 for each group at 3, 6, 8 days post infection. Data represented as mean ± SEM. *, **, **** denotes significance of *p* < 0.05, *p* < 0.01, *p* < 0.0001, respectively, as (B) determined by repeated measures two-way ANOVA corrected for multiple comparisons (Šídák). EF50, expiratory flow at 50% expired volume; PIF, peak inspiratory flow; PEF, peak expiratory flow; PAU, pause; PENH, enhanced pause.

**Table 5A.**
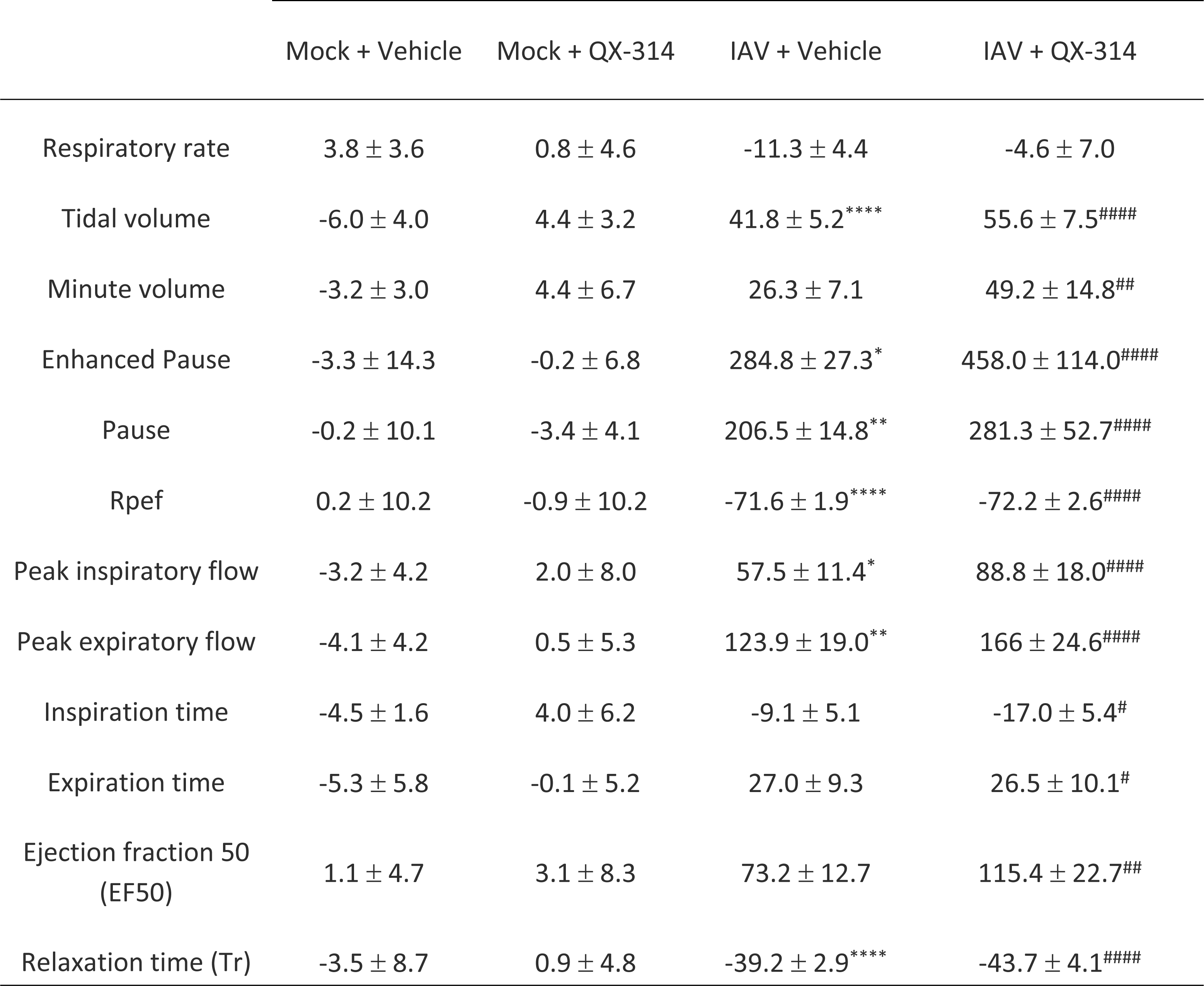
Whole body plethysmography at day 3 post-infection. % change from baseline measurements (Day 0, prior to infection).

**Table 5B.** Whole body plethysmography at day 6 post-infection. % change from baseline measurements (before infection).

|  | Mock + Vehicle | Mock + QX-314 | IAV + Vehicle | IAV + QX-314 |
| --- | --- | --- | --- | --- |
| Respiratory rate | 0.6 ± 3.0 | 2.2 ± 3.7 | -24.1 ± 2.3** | -19.6 ± 3.6## |
| Tidal volume | -2.2 ± 5.4 | 1.0 ± 1.6 | 56.3 ± 7.9**** | 68.2 ± 8.3#### |
| Minute volume | -1.2 ± 5.9 | 2.3 ± 4.8 | 20.1 ± 8.2 | 36.9 ± 10.7# |
| Enhanced Pause | 2.6 ± 13.2 | 2.8 ± 8.4 | 705.6 ± 64.0**** | 1006.6 ± 87.1####, ^ |
| Pause | -2.4 ± 9.6 | -1.4 ± 4.5 | 449.0 ± 35.0**** | 547.2 ± 39.2#### |
| Rpef | 7.0 ± 9.5 | 10.7 ± 11.5 | -77.2 ± 2.2**** | -77.7 ± 1.9#### |
| Peak inspiratory flow | 2.4 ± 6.8 | 0.2 ± 5.0 | 72.1 ± 14.6** | 94.9 ± 16.8#### |
| Peak expiratory flow | 6.7 ± 5.7 | 0.9 ± 4.4 | 190.5 ± 26.6**** | 255.0 ± 31.3#### |
| Inspiration time | -5.6 ± 6.1 | 2.8 ± 5.3 | -11.1 ± 4.3 | -14.9 ± 3.7 |
| Expiration time | 0.8 ± 4.7 | -1.9 ± 4.2 | 61.3 ± 6.0**** | 53.1 ± 11.0#### |
| Ejection fraction 50 (EF50) | 3.2 ± 5.0 | 4.6 ± 6.0 | 144.2 ± 26.8** | 230.1 ± 40.1####, ^ |
| Relaxation time (Tr) | 5.6 ± 6.7 | -1.5 ± 4.0 | -53.0 ± 2.6**** | -60.0 ± 3.0#### |

**Table 5C.**
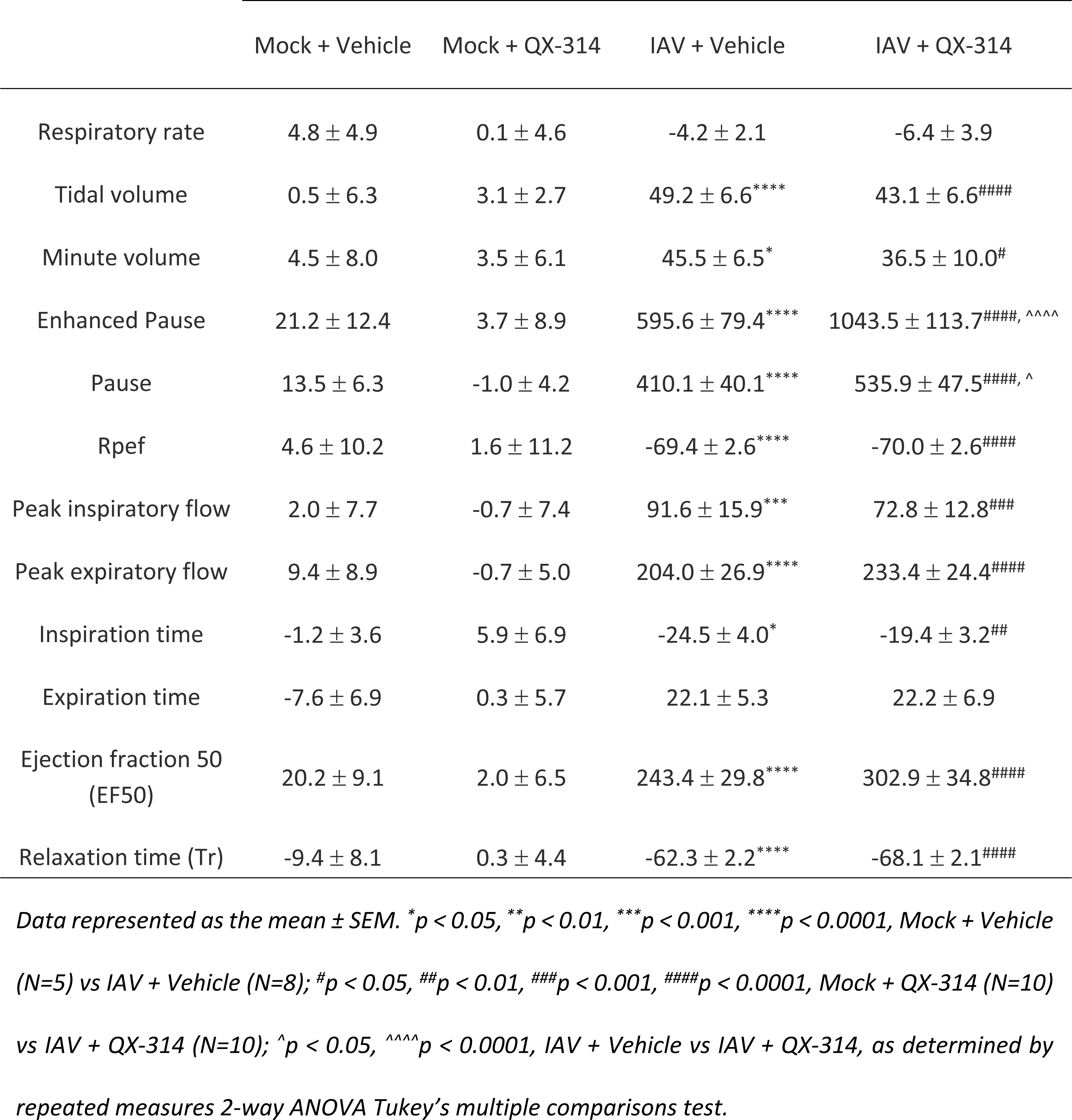
Whole body plethysmography at day 8 post-infection. % change from baseline measurements (before infection).

No differences were observed in proinflammatory lung cytokines, lung histopathology (Supp Fig 2C and D) or immune cells present in the BAL (Tables 6A-C) between mock infected QX-314 and vehicle mice. In IAV infected mice, across both QX-314 and vehicle groups, there was an obvious increase in proinflammatory cytokines in the lungs (IL-6, IL-1β, IFNγ, TNFα; Fig 3A), significant pathology present in the lungs (Fig 3B) and changes in the lung immune cell populations (Table 6A-C). We did observe a significant decrease in IL-1β (IAV + QX-314: 337.6 ± 40.8 pg/ ml, IAV + vehicle: 685.6 ± 159.2 pg/ ml, *p* = 0.0043; Fig 3A) and IFNγ (IAV + QX-314: 155.5 ± 18.2 pg/ ml, IAV + vehicle: 489.0 ± 114.4 pg/ ml, *p* = 0.0030; Fig 3A) and a small improvement in lung histopathology (Fig 3B) at day 8 post-infection in IAV infected QX-314 mice compared to IAV infected vehicle mice. In addition, interstitial macrophages were significantly more frequent at day 4 post-infection with a reduction in B-cells and CD8^+^ T-cells at day 8 post-infection in IAV infected QX-314 mice compared to their vehicle counterparts. These data suggest that QX-314 administration leads to an altered regulation of the IAV immune response in the lungs. However, the nature of this pulmonary neuroimmune regulation appears inconsistent with the worsening of clinical signs observed when vagal sensory neurons are disrupted, or their activity prevented.

**Figure 3.**
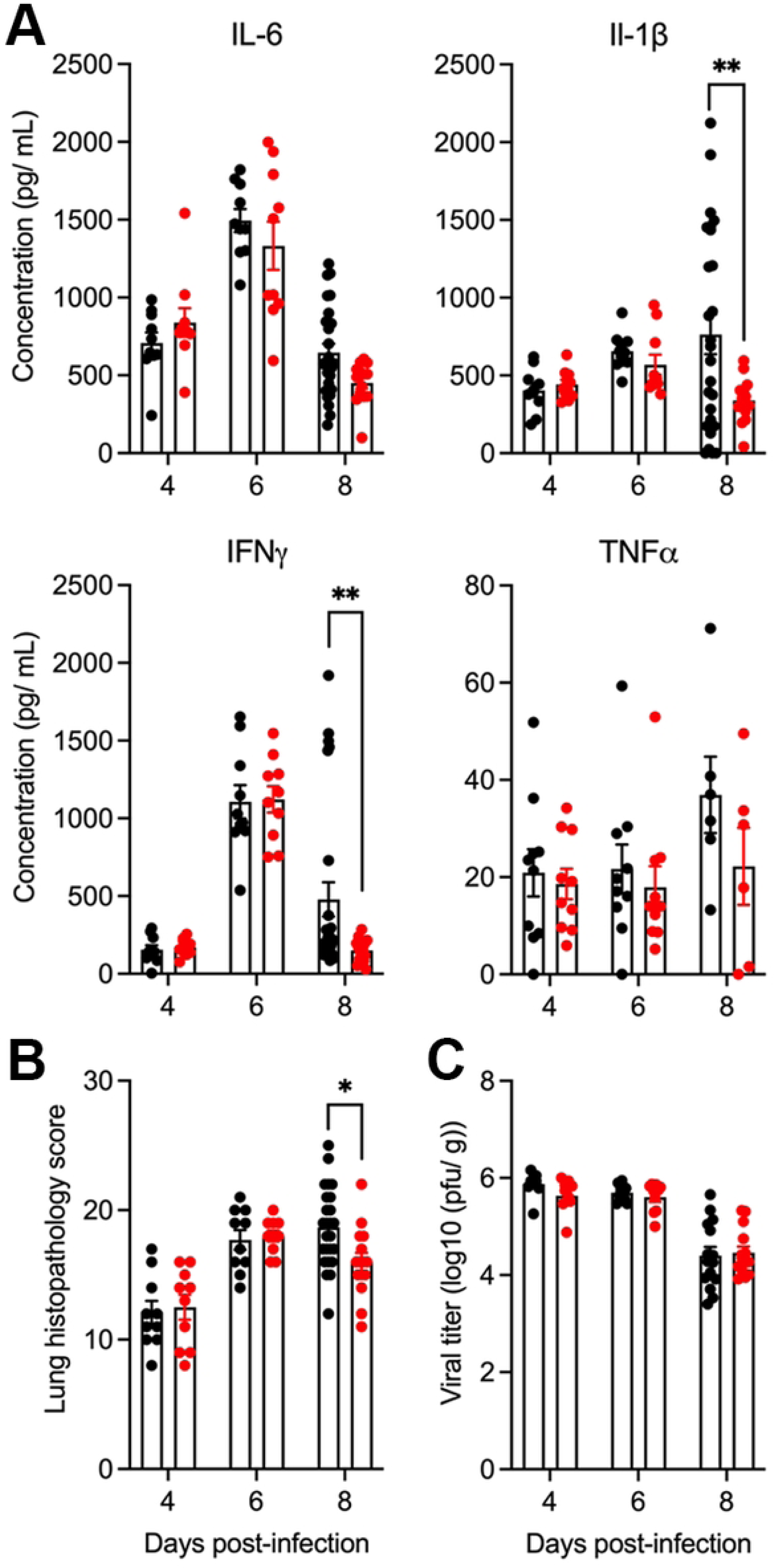
Pharmacological inhibition of pulmonary sensory neuron activity impacts IAV pathogenesis. Graphs depict group level (A) lung cytokines, (B) lung histopathological score, and (C) lung viral titers post infection with IAV following administration of QX-314 or vehicle. N = 10 for each group at 3, 6, 8 days post infection. Data represented as mean ± SEM. *, ** denotes significance of *p* < 0.05, *p* < 0.01, respectively, as (B) determined by repeated measures two-way ANOVA corrected for multiple comparisons (Šídák).

**Table 6A.**
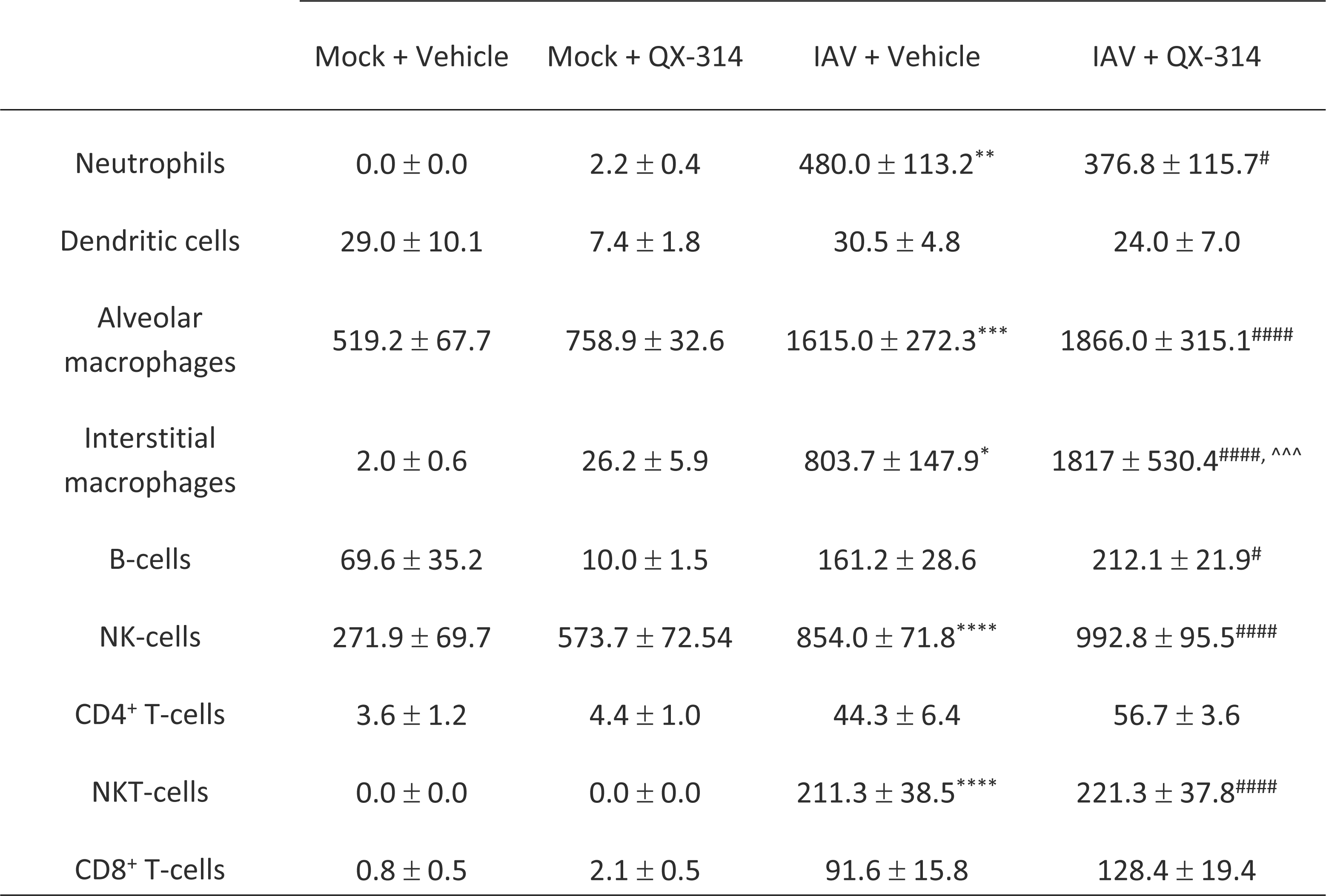
Flow cytometry of BALF at day 4 post-infection.

**Table 6B.** Flow cytometry of BALF at day 6 post-infection.

|  | Mock + Vehicle | Mock + QX-314 | IAV + Vehicle | IAV + QX-314 |
| --- | --- | --- | --- | --- |
| Neutrophils | 0.0 ± 0.0 | 3.7 ± 1.5 | 1289.0 ± 100.1**** | 1193.0 ± 185.4#### |
| Dendritic cells | 29.0 ± 10.1 | 10.0 ± 2.5 | 52.4 ± 7.7 | 34.5 ± 3.9 |
| Alveolar<br>macrophages | 519.2 ± 67.7 | 824.4 ± 117.4 | 1265.0 ± 108.4* | 1300.0 ± 127.5 |
| Interstitial<br>macrophages | 2.0 ± 0.6 | 4.7 ± 1.8 | 375.2 ± 36.6 | 436.0 ± 75.3 |
| B-cells | 69.6 ± 35.2 | 65.4 ± 10.8 | 516.4 ± 80.5 | 493.3 ± 33.2# |
| NK-cells | 271.9 ± 69.7 | 129.3 ± 19.2 | 197.5 ± 11.5 | 211.6 ± 22.4 |
| CD4<sup>+</sup> T-cells | 3.6 ± 1.2 | 4.0 ± 0.9 | 170.0 ± 34.2 | 121.3 ± 9.6 |
| NKT-cells | 0.0 ± 0.0 | 0.0 ± 0.0 | 151.5 ± 22.5*** | 133.9 ± 22.8### |
| CD8<sup>+</sup> T-cells | 0.8 ± 0.5 | 2.0 ± 0.7 | 259.5 ± 70.2 | 379.1 ± 92.9 |

**Table 6C.** Flow cytometry of BALF at day 8 post-infection.

|  | Mock + Vehicle | Mock + QX-314 | IAV + Vehicle | IAV + QX-314 |
| --- | --- | --- | --- | --- |
| Neutrophils | 0.0 ± 0.0 | 0.0 ± 0.0 | 488.6 ± 63.8* | 295.4 ± 77.3 |
| Dendritic cells | 29.0 ± 10.1 | 48.1 ± 14.3 | 440.0 ± 116.0**** | 256.4 ± 74.5#### |
| Alveolar macrophages | 519.2 ± 67.7 | 670.9 ± 121.9 | 850.9 ± 97.1 | 880.4 ± 208.4 |
| Interstitial macrophages | 2.0 ± 0.6 | 1.2 ± 0.6 | 173.2 ± 34.3 | 124.5 ± 47.0 |
| B-cells | 69.6 ± 35.2 | 77.6 ± 22.4 | 849.0 ± 111.3**** | 638.3 ± 155.6####, ^ |
| NK-cells | 271.9 ± 69.7 | 541.0 ± 155.4 | 527.1 ± 74.2 | 384.1 ± 102.2 |
| CD4<sup>+</sup> T-cells | 3.6 ± 1.2 | 3.2 ± 1.1 | 1534.0 ± 141.4**** | 1528.0 ± 457.5#### |
| NKT-cells | 0.0 ± 0.0 | 0.0 ± 0.0 | 46.5 ± 13.0 | 24.8 ± 7.3 |
| CD8<sup>+</sup> T-cells | 0.8 ± 0.5 | 1.4 ± 0.5 | 2974.0 ± 447.3**** | 1499.0 ± 456.8####,<br>^ ^ ^ ^ |

*Data represented as the mean ± SEM. \*p < 0.05, \*\*p < 0.01, \*\*\*p < 0.001, \*\*\*\*p < 0.0001, Mock + Vehicle* *(N=5) vs IAV + Vehicle (N=5-10); #p < 0.05, ###p < 0.001, ####p < 0.0001, Mock + QX-314 (N=5-10) vs IAV* *+ QX-314 (N=5-10); ^p < 0.05, ^^p < 0.001, ^^p < 0.0001, IAV + Vehicle vs IAV + QX-314, as determined* *by repeated measures 2-way ANOVA multiple comparisons test.*

### Pharmacological inhibition of pulmonary sensory neurons impacts IAV neuropathology

We have previously shown that severe pulmonary IAV infections in mice are accompanied by a neuroinflammatory phenotype in the vagal sensory ganglia [19, 20]. To investigate whether peripheral sensory neuron activation in the lungs is involved in the development of vagal neuroinflammation, we compared the expression of inflammatory genes and inflammatory cell densities in the vagal sensory ganglia of IAV infected mice administered via inhalation from day 3 post IAV infection with either vehicle or QX-314 (Fig 2A). Consistent with our previous studies, in vehicle administered mice, IAV compared to mock infection resulted in the increase of expression of several inflammatory related genes within the vagal sensory ganglia, *Tmem173* (fold change: day 4, 0.3 ± 0.5, *p* = 0.021; day 8, 2.7 ± 0.3, *p* < 0.0001), *Il1b* (fold change: day 4, 140.4 ± 2.2, *p* < 0.0001; day 8, 6.3 ± 1.5, *p* < 0.0001), *Irf9* (fold change: day 4, 2.0 ± 0.2, *p* < 0.0001; day 8, 2.4 ± 0.1, *p* < 0.0001)*, and Isg15* (fold change: day 4, 0.8 ± 0.04, *p* = 0.24; day 8, 12.2 ± 3.4, *p* = 0.005). Inflammatory gene changes were accompanied by a significant increase in the number of ganglia MHC II+ immune cells of IAV compared to mock infected mice at day 6 and day 8 post-infection (IAV + vehicle: day 6, 58.8 ± 2.1 % and day 8, 73.6 ± 2.4 %; Mock + vehicle: 38.9 ± 1.9 % MHC II+ cells relative to total neurons; *p* = 0.0011 (D6) and *p* < 0.0001 (D8)).

QX-314 administration resulted in an altered neuropeptide and inflammatory gene profile in the vagal sensory ganglia. In IAV infected QX-314 administered mice, we observed a significant increase in the expression of genes encoding the neuropeptides CGRP (*Calca*) and Substance P (*Tac1*) at both days 4 and 8 post infection compared to IAV vehicle mice. We also observed a further upregulation of genes associated with host defense against pathogens, *Tmem173* and *Il1b*, and a decrease in genes associated with an interferon antiviral response, *Irf9* and *Isg15* (Fig 4A). The H1N1 subtype of IAV is typically considered non-neurotropic, yet as we have previously reported [19] we detected viral mRNA in the vagal sensory ganglia at days 4 and 6 post infection of mice administered vehicle. Intriguingly, mice administered QX-314 demonstrated lower ganglia viral mRNA levels, suggestive of lower infection rates or improved viral clearance (Fig 4B). Mock infected mice administered either vehicle or QX-314 showed no difference in the number of MHC II+ immune cells and CGRP-expressing neurons within their vagal sensory ganglia (Supp Fig 3A and B). However, IAV infection resulted in a further increase in the number of MHC II+ cells with mice that were administered QX-314 displaying a significantly higher number of MHC II+ immune cells at days 6 and 8 post-infection compared to IAV infected vehicle mice (Fig 4C). We also observed a significant increase in the overall number of CGRP-expressing neurons within the vagal sensory ganglia at days 4 and 6 post-infection in QX-314 mice compared to their vehicle counterparts (Fig 4D).

**Figure 4.**
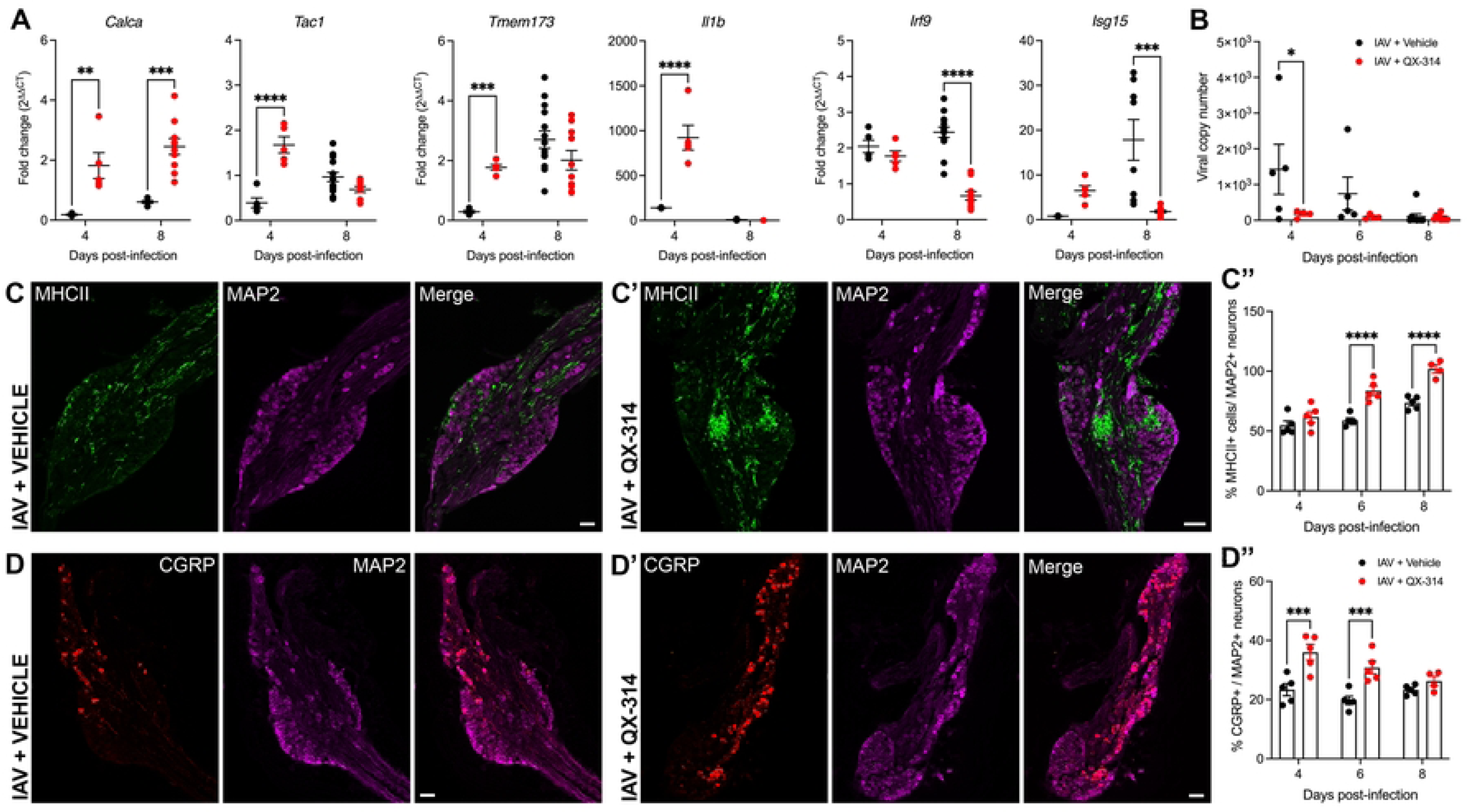
Pharmacological inhibition of pulmonary sensory neuron activity during IAV infection results in a hyperinflammatory state within the vagal sensory ganglia. Graphs depict group level qPCR analysis of (A) neuropeptide and host defense/ proinflammatory associated genes, and (B) viral mRNA levels present in the vagal sensory ganglia during IAV infection in mice that received either vehicle (black circles) or QX-314 (red circles). Data expressed as fold change in expression values relative to matched mock group. Representative immunofluorescence images of the vagal sensory ganglia at day 6 post-infection demonstrating an increase in the number of MHC II+ immune cells in mice administered either (C) vehicle or (C’) QX-314 (MHC II, green; sensory neurons immunolabeled with MAP2, purple). The bar graph (C”) shows the percentage increase of MHC II+ cells per total number of MAP2+ neurons in the vagal sensory ganglia of vehicle (black circles) and QX-314 (red circles) IAV infected mice. Representative immunofluorescence images of the vagal sensory ganglia demonstrating an increase in the number of CGRP-expressing neurons in IAV infected mice that received (D’) QX-314 compared to mice receiving vehicle (D). The bar graph (D”) shows the percentage increase of CGRP-expressing neurons (red) per the total number of MAP2 neurons (purple) in the in the vagal sensory ganglia of vehicle (black circles) and QX-314 (red circles) IAV infected mice. Data represented as mean ± SEM. *, **, ***, **** denotes significance of *p* < 0.05, *p* < 0.01, *p* < 0.001, *p* < 0.0001, respectively, as (A, B, C”, D”) determined by repeated measures two-way ANOVA or mixed-effects analysis corrected for multiple comparisons (Šídák). Scale bar represents 100μm.

## Discussion

There is a growing recognition that neuroimmune crosstalk contributes to the pathogenesis and clinical presentation in many diseases, potentially representing a novel therapeutic checkpoint for improved disease management. The present study advances this proposition by demonstrating that vagal neural processes contribute to the regulation of inflammation and morbidity during IAV infection. Denervating the lung of its vagal sensory neural innervation slightly increased IAV pathology, whereas pharmacologically preventing pulmonary sensory neuron activation during IAV infection marginally improved overall lung inflammation and pathology, altered the accompanying vagal sensory ganglia neuroinflammation immune response and worsened overall disease. Collectively, these data build on our understanding of neuroimmune interactions and suggest complex roles for the nervous system in pulmonary viral disease.

Vagal pathways have been shown to regulate pulmonary inflammation in dichotomous ways. For example, in a murine model of allergic asthma the inflammatory cytokine IL-5 was shown to trigger the release of the neuropeptide vasoactive intestinal protein (VIP) from a subset of pulmonary sensory neurons which, in turn, perpetuated inflammation by acting on CD4^+^ T cells and innate lymphoid cells, increasing the production of TH2 cytokines associated with driving asthmatic conditions. Ablating these pulmonary sensory neurons reduced asthma severity [21]. The same authors showed that blocking the activity of pulmonary sensory neurons and the subsequent release of substance P during allergic asthma was sufficient to reduce allergy-induced goblet cell hyperplasia and hypersecretion of Muc5AC [22], a mucin that contributes to airway hyperactivity [35]. However, not all effects of vagal sensory neuron activation are detrimental. CGRP released from vagal lung sensory neurons during pulmonary staphylococcal infection mediated immunosuppression and reduced intracellular killing of *S. aureus* by neutrophils. CGRP administration increased the bacterial load in the lungs of infected mice and worsened the clinical signs of pneumonia [23]. These observations suggest that pulmonary sensory neurons may play complex and multifaceted roles in respiratory disease, perhaps specific to the underlying pathogen or disease pathology.

In the present study, both unilateral cervical vagotomy and genetic or pharmacological approaches to ablate or inhibit pulmonary sensory neurons provided evidence for vagal controls over IAV-associated pulmonary inflammation. In accordance with our findings, Gao and colleagues reported that vagotomy reduced lung anti-inflammatory responses during a PR8 infection in mice. This effect was presumably due to a disruption of vagal contribution to the systemic anti-inflammatory reflex which is governed by autonomic innervation to splenic and other abdominal immune tissues [36]. Similar effects of vagotomy have also been reported during non-pathogenic systemic lipopolysaccharide exposure [37]. By contrast vagotomy has been shown to reduce pulmonary inflammation associated with lung fibrosis [38]. These findings suggest that vagal mechanisms may contribute to inflammatory regulation via both local tissue and systemic pathways, perhaps complicating interpretation of such studies. Indeed, vagotomy non-specifically disrupts all sensory and motor innervation to the tissues below that vagotomy level, and therefore we cannot presume that the effects observed in our vagotomy studies are specifically related to either sensory pathways or the innervation to the lungs. However, it is notable that systemic cytokines released during IAV infection were not altered by vagotomy, which may argue against a substantive disruption of the systemic anti-inflammatory reflex. Furthermore, anti-inflammatory effects were observed following QX-314 inhalation treatment in mice exposed to IAV, which presumably preserves the efferent pathways regulating the systemic anti-inflammatory reflex.

The mechanisms involved in vagal sensory regulation of IAV pulmonary inflammation are not clear but could involve local neuropeptide-dependent processes. We and others have previously observed an increase in neuropeptide gene and protein expression markers in vagal sensory neurons following pulmonary viral infections [19] and administration of QX-314 in IAV infected animals elicited a further increase in vagal ganglia neuropeptide content in the present study. Regardless of the mechanism, it is intriguing that the anti-inflammatory effects of vagal sensory nerve manipulations in the present study did not impact viral load in the lungs. Nevertheless, influenza virus infected animals consistently presented with greater weight loss and worsened clinical scores following vagotomy, sensory neuron ablation or inhibition, seemingly incongruent with the moderate pulmonary inflammatory effects and unchanged infectious burden. These findings suggest that alternative mechanisms are involved in sensory neuron-dependent impacts on IAV morbidity.

In our previous study, we characterized transcriptomic changes and an increase in inflammatory cells in the vagal sensory ganglia following pulmonary infection with IAV, consistent with IAV-induced neuroinflammation [19]. Upregulated genes were mostly related to pathogen host defense and anti-viral responses, typically downstream of interferon signalling, suggesting that pulmonary sensory neurons are influenced by the local antiviral inflammatory environment in the lungs during IAV infection. Here, we replicated these findings, and extend our prior observations by providing further evidence that vagal ganglia neuroinflammation is likely regulated through mechanisms that involve sensory neuron activation in the lungs. Thus, the expression of the interferon-related signalling genes *Irf9* and *Isg15* in the vagal sensory ganglia was reduced in animals receiving QX-314, conceivably related to the concomitant reduction in lung IFNγ, known to activate the heterotrimer Stat1:Stat2:Irf9 complex (interferon stimulating gene factor 3; ISGF3) to enhance transcription of interferon stimulated genes (ISGs) and pro-inflammatory cytokines [39, 40].

Intriguingly, however, the effects of QX-314 on vagal ganglia neuroinflammation were not as predicted, as inflammatory cell numbers in the vagal ganglia were significantly increased in IAV-infected animals receiving QX-314 and several other pro-inflammatory genes were upregulated, including the inflammasome associated genes *Il1b* and *Tmem173* (encodes for stimulator of interferon genes; STING) [41, 42]. The inflammasome in neurons is likely a defense mechanism against viral replication [43, 44] and whilst H1N1 IAV is not typically considered a neurotropic virus, previous studies have reported the presence of viral RNA in neural tissue such as the vagal ganglia and pulmonary sensory neurons following pulmonary infection [19]. It is interesting to speculate that diminished recruitment of interferon-dependent signalling pathways following QX-314 administration may lead to enhanced activation of sensory neuron inflammasome signalling to combat IAV reaching and replicating within the vagal ganglia. Indeed, the viral copy number of IAV was decreased in the vagal sensory ganglia of animals that received QX-314. The potentiation of the increased number of ganglia immune cells (MHC II expressing) and upregulated neuropeptide expression could also be part of an increased immune response against IAV neuronal invasion. However, we cannot rule out these findings reflect an alternative response to, or cause of, the transcriptomic changes through currently unknown mechanisms. Regardless, the altered neuroinflammatory profile following inhibition of the activity of pulmonary sensory neurons with QX-314 may result in broad neurological impacts and underpin the altered clinical presentation evident in these animals.

In conclusion, these data show that the pulmonary vagal sensory neurons play a role in the regulation of immune responses in the pulmonary system during IAV infection. Surgical, genetic, and pharmacological interventions targeting the vagus nerve and sensory neurons had modest effects on lung inflammation and viral pathogenesis but worsened the clinical presentation perhaps due to a dysregulated vagal neuroinflammation. The exact mechanisms by which this occurs remains to be elucidated. Whether the neuroinflammatory impacts of IAV pathogenesis extend beyond the vagus nerve into the central nervous system and circuits regulated by pulmonary vagal sensory neurons warrants further investigation. Conceivably, central mechanisms could contribute to the myriad of disease behaviours, such as malaise, myalgia, and cough that accompany IAV and other severe lung infections [45, 46]. Targeting specific vagal sensory pathways could thus be a potential therapeutic approach to improve IAV disease outcomes.

## Materials and Methods

### Virus details

Viral stocks of Influenza Auckland/ 1/ 2009 (Auck/ 09; H1N1) and Puerto Rico/ 8/ 34 (PR8, H1N1) were propagated as previously described in embryonated chicken eggs, approved by the University of Queensland Animal Ethics Committee (AE000089). Viral titres of IAV were determined by plaque assay on Madin-Darby Canine Kidney (MDCK) cells, as previously described [47]. We note the use of the PR8 strain of H1N1 with the vagotomy experiments and the Auck/ 09 H1N1 strain with the experiments involving pharmacological inhibition of neural activity. An initial characterization revealed no differences of IAV pathogenesis between the two viral strains (Supp Fig 4) and we also observed a comparable neuroinflammatory profile within the vagal sensory ganglia (Fig 4 and [19]).

### Murine model

#### Mouse strains

This study complies with the *Australian Code for the Care and Use of Animals for Scientific Purposes* from the National Health and Medical Research Council of Australia. All procedures were approved by the Animal Ethics Committee of both The University of Queensland, Australia and the University of Melbourne, Australia (approval number 071/17 and 20986, respectively). All steps were taken to minimize the animals’ pain and suffering, as mandated in the Australian Code. Mice were housed in temperature controlled (21 °C), individually ventilated cages on a 12-hour light/ dark cycle, with access to pelletized food and water ad libitum in groups up to five per cage. Pathogen-free wildtype C57BL/6JArc mice (8-10 weeks of age, male) were obtained from the Animal Resources Centre (ARC; Western Australia, Australia) and used for all experiments involving surgical denervation, retrograde neuronal tracing and QX-314 treatment. Heterozygous Phox2b-Cre mice (Stock No: 016223; B6(Cg)-Tg (Phox2b-Cre)3Jke/J) were initially purchased from the Jackson Laboratory, with a colony bred and maintained at the University of Melbourne in accredited facilities. Phox2b mice expressing Cre (Phox2b-Cre^+^) and littermate controls (Phox2b-Cre^-^) used for experimentation (8-12 weeks of age, male and female) were generated by crossing male heterozygous Cre mice to female wildtype C57BL/ 6 mice. Mice were genotyped for Cre as per genotyping guidelines on the Jackson Laboratory website.

#### Surgical Procedures

A unilateral vagotomy was performed to partially disrupt nerve supply to the lungs. Mice were anesthetized with gaseous isoflurane (4% induction, 1.25-1.5% maintenance) and once their withdrawal and palpebral reflexes had disappeared a midline incision was made on the ventral surface of their neck to expose the right or left mid-cervical vagus nerve. Care was taken to carefully dissect each vagus nerve from the carotid artery and cervical sympathetic nerve. To avoid reinnervation approximately 7mm segment of nerve was removed, the wound was sutured, and mice were allowed to recover for at 10-14 days during which time body weight was monitored and post-operative analgesia was administered for 48 hours (Meloxicam, 5mg/ kg and buprenorphine 0.05mg/ kg s.c). A control sham group receiving the surgical procedure (everything described above omitting the removal of the vagus nerve) were also included. All mice, on average had reached pre-surgical body weight on the day of IAV infection (range, mean ± SEM of % body weight change from original; sham: 99.2 – 110.6 %, 105.2 ± 0.5 %; vagotomy: 97.9 – 107.8 %, 102.2 ± 0.4 %).

To determine the laterality of vagal sensory innervation to the right and left lung lobes we performed a series of experiments utilizing the retrograde viral tracer, AAV CAG-tdTomato (AAVrg^TdT^; Addgene 59462-AAVrg; 3μl of 4.6 x 10^12^ GC/ ml). Mice were split into 3 groups whereby group 1 had both vagi intact, group 2 had left vagus nerve intact (right vagotomy), and group 3 had right vagus nerve intact (left vagotomy). Within each group mice were further split into 2 subgroups whereby they received an injection into either the left or right lung lobes. Briefly, mice were injected from the posterior side and through the seventh intercostal space as previously described [15]. Skin and external intercostal musculature was retracted, the lung visualized underneath the intact internal intercostals and, using a 30 G needle attached to a 10 μl gastight Hamilton syringe, 3 μl of AAVrg^TdT^ virus was injected into the lung lobe. The needle was left in place for 2 min to limit the spread of AAV onto surrounding tissues. The incisions were sutured, and animals were allowed to recover for 21 days during which body weight was monitored daily and post-operative analgesics administered (see above).

To genetically ablate lung nodose sensory neurons, Phox2B-Cre mice received an injection of 6 μl AAVretro CAG-mCherry-FLEX-DTA (AAVrg^mCh-FLEX-DTA^; 4.85 x 10^11^ vg/ ml) diluted in 10 μl of sterile saline, slowly (over 2 minutes) into the tracheal lumen ∼5-6 cartilage rings below the larynx via a 10 μl Hamilton syringe connected to a 30 G needle, the needle was left in place for 30 s to prevent backflow following which the incision was sutured. Animals were allowed to recover for 28 days during which body weight was monitored daily and post-operative analgesics administered (see above).

#### IAV infection

Mice were infected intranasally (50 μl) with 50 PFU A/Puerto Rico/8/1934 H1N1 (PR8) or 5.5 x10^3^ PFU of A/Auckland/1/2009 H1N1 (Auck/09) suspended in phosphate buffered saline (PBS, pH 7.4, Thermo Fisher, Australia) under isoflurane-induced anaesthesia (1-3 L/ min) with PBS being used for mock infections. Over the course of disease, mice were monitored at the same time every day for body weight changes and clinical signs associated with disease progression. The scoring criteria included in the clinical signs assessment include:

1. Percentage weight loss calculated from pre-infection weight (no change OR 1-20 % loss of body weight OR > 20 % loss of body weight).
2. General condition, which included appearance of fur (shiny and unruffled OR slightly ruffled OR ruffled and matte), eye condition (clear, clean, and open OR unclean, evidence of discharge, semi-closed/ closed), posture (normal OR slightly hunched OR moderately hunched OR severely hunched), and clinical complications (paralysis, tremor AND/ OR vocalizations, breathing sounds AND/ OR animal is appearing cold to touch, decrease in body temperature).
3. Motility (spontaneous normal behavior with social contacts OR spontaneous but reduced OR moderately reduced behavior OR motility only after stimulation OR isolation, coordination disorders, not responding to stimulation).
4. Respiration (breathing normal OR breathing slightly changed i.e., hyper/ hypoventilation, labored/ shallow breathing/ gasping, up to 10 % change OR breathing moderately altered, 10 – 30 % change OR breathing strongly altered, 30 – 50 % change).

#### QX-314 treatment

Mice were nebulized with lidocaine N-ethyl bromide (N-(2,6-Dimethylphenylcarbamoylmethyl) triethylammonium bromide (QX-314; Sigma Aldrich, Australia) dissolved in sterile saline (300µM) or vehicle (sterile saline), twice daily (12 hrs apart) beginning at three days post IAV infection. This consisted of mice (groups of five) placed in a Buxco small animal whole body plethysmography chamber (diameter, 23cm; height, 12 cm; volume, 4985 cm^3^) and exposed to aerosolized QX-314 or vehicle (Buxco Aerosol Distribution Unit 5 LPM, Aerogen nebulizer unit AG-AL1000 with filter cap of 3.1 µm particle size; Data Sciences International) over 20 minutes, at an air flow velocity of 2.5 L/ min and at 50% nebulizer duty cycle. The inhaled dose of QX-314 for each animal was, on average 0.04 mg/ kg. This value was calculated using this algorithm, in accordance with [48]: Inhaled dose (mg/kg) = [C (mg/ L) x RMV (L/ min) x D (min)]/ BW (kg), where C is the concentration of drug in air inhaled, RMV is respiratory minute volume, D is the of exposure in minutes, BW is bodyweight in kg, with RMV (in L/ min) calculated as 0.608 x BW (kg)^0.852^.

#### Whole-body plethysmography recordings

Respiratory parameters were measured in mice (QX-314 experimental group) using calibrated 4-chamber unrestrained whole-body plethysmography (WBP, Buxco, Wilmington, USA). Mice were acclimatized to the plethysmography chambers (30 minutes per day), each day for 3 days prior to experimentation. Respiratory frequency (f, breaths min ^-1^), tidal volume (TV, ml kg ^-1^), minute volume (MV, ml min ^-1^), inspiratory and expiratory time (Ti and Te, sec), estimated peak inspiratory and expiratory flow (PIF and PEF, ml sec ^-1^), pause and enhanced pause (PAU and Penh, arbitrary unit of measure), expiratory flow at 50% expired volume (EF50, ml sec -1), and relaxation time (Tr, sec) were automatically sampled from the calibrated Buxco flow trace, corrected for animal body weight and ambient chamber temperature, and calculated in real time by the plethysmography software (Buxco Finepointe Software Version 2.1.0.9). Real time data collected every 2 sec were used to calculate the average per 20 min for each animal at baseline (day 0, prior to infection) and days 4, 6, 8 post-infection. Data was normalized to baseline values for each animal. Normalization was performed to remove confounding factors such as inter-animal variation in baseline parameters.

### Tissue sampling and analyses

All tissue samples were obtained from mice that were euthanized post-infection with sodium pentobarbital (100 mg/ ml) and exsanguinated unless otherwise stated.

#### Viral plaque assay

Lung lobes were homogenized in DMEM (ThermoFisher Scientific, Australia), clarified by centrifugation, and stored at -80°C until analysis. Viral titers in clarified tissue homogenate samples were determined by plaque assay, as described previously [49].

#### Cytokine measurements

Cytokine levels in clarified tissue lung homogenates were determined by enzyme linked immunosorbent assay (ELISA) according to the manufacturer’s instructions (BD Biosciences, USA) and read with CLARIOstar Plus (BMG labtech) microplate reader. Serum cytokine levels were determined using the Anti-virus Response panel of LEGENDplex^TM^ Multiplex Assays (BD Biosciences, USA) according to the manufacturer’s instructions, measured using the Cytoflex S flow cytometer (Beckman Coulter, USA) and analysed using LEGENDplex Qognit software (BD Bioscience, USA).

#### Broncho-alveolar lavage fluid (BALF) immune cell populations

BALF samples were clarified by centrifugation with red blood cells (RBC) lysed in RBC lysis buffer (0.15 M NH_4_Cl, 17 nM Tris-HCl at pH 7.2, sterile filtered) for 1 min and cell pellet resuspended in 0.1M PBS. Samples were treated with BD Fc Block (Rat Anti-mouse CD16/CD32; BD Biosciences, USA) for 20 min at 4 °C, followed by incubation in the following fluorophore conjugated anti-mouse antibody combinations for 20 min at 4 °C: FITC NK1.1 (clone PK136), PerCP-Cy5.5 anti-CD8a (clone 53-6.7), PE/ Cy7 anti-CD3 (clone 53-6.7) (BD Biosciences, USA); FITC anti-CD11b (clone M1/ 70), PE anti-CD4 (clone GK1.5), PerCP-Cy5.5 anti-GR-1 (clone RB6-8C5), APC anti-F4/ 80 (clone BM8), APC anti-B220 (clone RA3-6B2) (Biolegend, USA); PE/Cy7 anti-CD11c (clone N418) (eBioscience, USA) and eFluor780 fixable viability dye (ThermoFisher Scientific, Australia). This was followed by fixation using BD Cytofix Fixation buffer (BD Biosciences, USA) to preserve immunofluorescent staining until analysed. Samples were analysed using the Cytoflex LX flow cytometer (Beckman Coulter, USA) and data analysed with FlowJo_v10 software. Immune cell populations were classified as followed: alveolar macrophages (F4/80^+^ CD11b^int/low^ CD11c^high^), interstitial macrophage (F4/80^+^ CD11b^high^ CD11c^int^), neutrophils (F4/80^-^ GR-1^high^ CD11b^high^), dendritic cells (F4/80^-^ GR-1^-^ CD11b^high^ CD11c^high^), CD4^+^ T cells (CD3^+^ CD4^+^), CD8^+^ T cells (CD3^+^ CD8^+^), NK cells (CD3^-^ NK1.1^+^), NKT cells (CD3^+^ CD4^-^ CD8^-^ NK1.1^+^). See Supp Fig 4 for gating strategy.

#### Pulmonary histology

Lung lobes were collected and fixed in 10 % formalin for a minimum of 24 h then transferred to 70% ethanol for processing by the Core Histology Facility, Translational Research Institute (Woolloongabba, Queensland, Australia). Lung samples were embedded in paraffin wax prior to sectioning at 7 µm using a Hyrax M25 Rotary Microtome (Leica Biosystem, Germany). Sections were stained with hematoxylin and ethanol-based eosin and assessed by a veterinary pathologist who was blinded to the study designs. Each sample was scored for vascular changes, bronchitis/ bronchiolitis, interstitial inflammation, alveolar inflammation, pneumocyte hypertrophy and pleuritis in a semi-quantitative manner on a scale of 0-5 where 0 = no change, 1= minimal change, 2 = mild change, 3 = moderate change, 4 = severe change in < 50 % of lung lobe and 5 = severe change in > 50 % of lung lobe.

#### Vagal sensory ganglia RNA isolation and quantitative PCR

Vagal sensory ganglia analyses were performed on mice as part of the QX-314 experiments (see above). Mice were euthanized at day 4, 6, 8 post-infection and bilateral vagal sensory ganglia freshly harvested, placed in 250 μl TRIzol reagent (Qiagen, Australia), snap frozen and stored at -80 °C until further processing. Thawed samples were homogenized and RNA was extracted and concentrated using the RNeasy MiniElute Cleanup kit (Qiagen, Australia) according to the manufacturer’s instructions. cDNA was generated using the high-capacity cDNA reverse transcription kit (Life technologies) and random primers (host RNA) according to the manufacturer’s instructions. Real-time PCR was performed on generated cDNA with SYBR Green using QuantStudio 6 Real-Time PCR (ThermoFisher Scientific, Australia). Forward and reverse prime sequences for each gene of interest are shown in Table 7. Gene expression was normalized relative to Hypoxanthine phosphoribosyl transferase (*Hprt*) and β-actin (*Actb*) expression. Fold change was calculated based on the 2^ΔΔCT^ method. Viral copy number was determined using the A/Puerto Rico/8/1934 H1N1 virus matrix gene cloned into the pHW2000 plasmid as previously described [50]. Primers used for IAV quantification, forward primer: AAGACCAATCTTGTCACCTCTGA, reverse primer: TCCTCGCTCACTGGGCA.

**Table 7.**
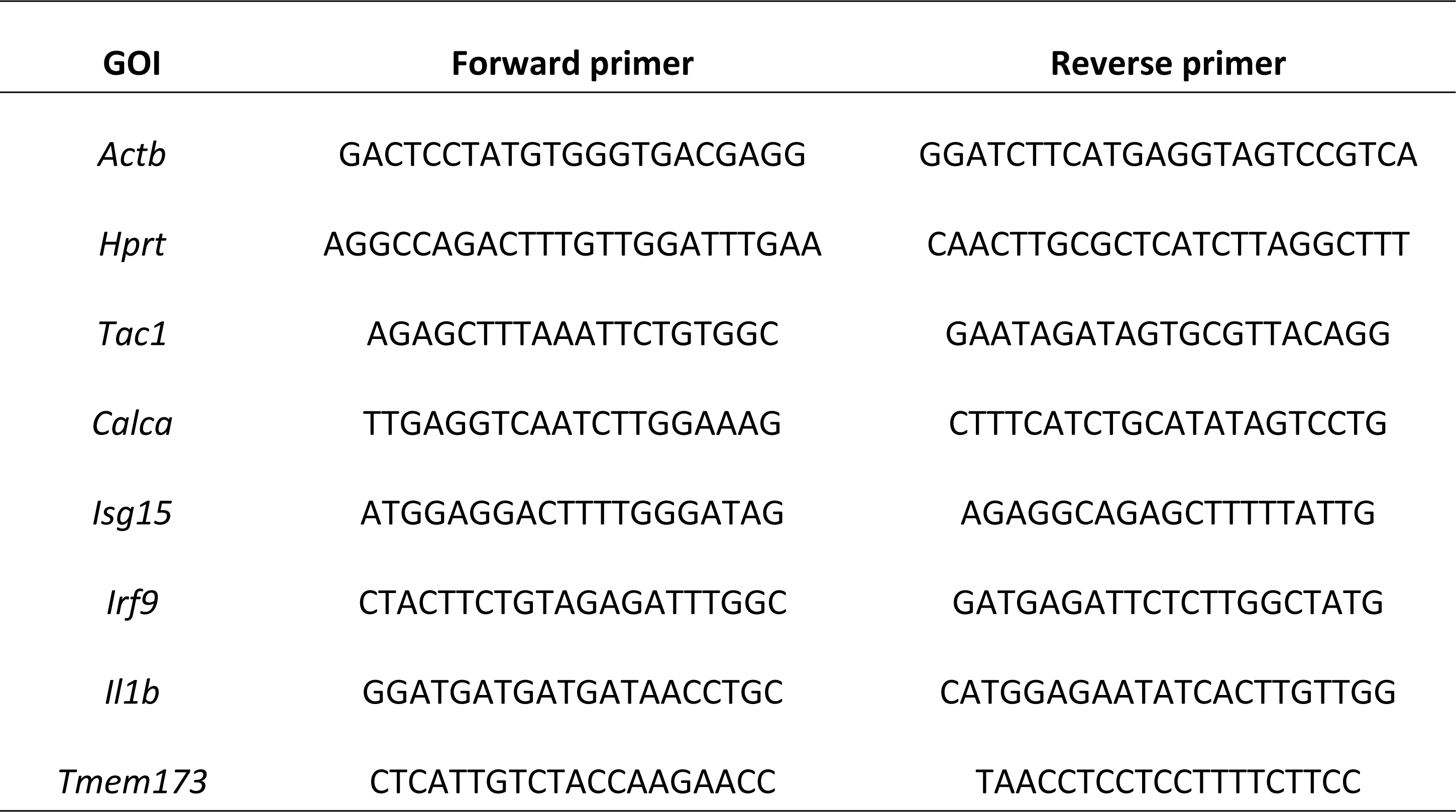
Primers used for vagal sensory ganglia qPCR.

#### Immunofluorescence assay of vagal sensory ganglia

A subset of mice from the QX-314 experiments and Phox2B-Cre mice were euthanized with (100 mg/ kg, i.p.) sodium pentobarbital and transcardially perfused with PBS; 0.1 M, pH 7.45, followed by 4 % PFA; pH 7.45. Vagal sensory ganglia and brainstems were removed and post-fixed in 4 % PFA (2 hours) then cryoprotected in 30 % sucrose (w/v PBS) overnight before freezing in Optimal Cutting Temperature compound (OCT, TissueTek). Twelve µm sections were collected on a cryostat (-20 °C), sequentially over 4 slides (Superfrost Plus) for each pair of vagal ganglia from one animal, dried at room temperature for 2 hrs and stored at -80 °C. 50 µm cryostat cut sections of Phox2B-Cre brainstems encompassing the medullary region were collected serially into 0.1M PBS and processed for immunohistochemistry that same day. All sections were rinsed in PBS then blocked in 10% donkey serum in 0.1 M PBS for 1 hr, before incubation for 24 hours in the primary antibody of interest: Rat anti I-A/ I-E, 1:1000, Biolegend, 107601; Chicken anti-microtubule associated protein 2 (MAP2), 1:1000, Abcam, Ab5392; Rabbit anti-calcitonin gene-related peptide (CGRP), 1:3000, Immunostar, 24112; Rabbit anti-dsRed, 1:1000, Clontech, 632496. All antibodies were diluted in antibody diluent (2% donkey serum and 0.3% Triton X-100 in PBS). After the required incubation, sections were washed three times with PBS for 20 mins each then incubated for one hour in appropriate fluorescently-conjugated secondary antibodies (1:500, ThermoFisher Australia). Brainstem sections were kept in order and mounted onto gelatin-coated slides. All slides were coverslipped with an antifade mounting media (ProLong Gold, ThermoFisher Scientific).

Sections were visualized on a fluorescent microscope (Leica DM6 B) with high resolution images taken with Leica DFC7000 T camera (x100) at the same exposure for each protein of interest across each sample for offline analysis. For vagal sensory ganglia, counts of inflammatory cells, neurons with MAP2 and CGRP or mCherry positive neurons were performed offline on all sections. Brainstem sections from Phox2B-Cre mice were viewed for the presence of mCherry positive fibers. High resolution digital images taken at 200x magnification were imported into Adobe Photoshop (version 20.0.4) and optimized (minimally) for brightness, contrast, and size for preparation of representative photomicrographs.

### Statistical analysis

Data are expressed as the mean ± SEM unless stated differently. Statistical analyses were performed using GraphPad Prism, version 8.3.1 for Windows (GraphPad Software, CA, USA). Normality of test groups were determined using Shapiro-Wilk’s test. Statistical significance for normally distributed data was determined using a two-tailed, unpaired students t-test (between two groups). A Mann-Whitney test was used for data that was not normally distributed. If one or more repeatedly measured variables were considered, uni- or multi-factorial repeated-measures ANOVA was performed. Statistical significance was set at *p* ≤ 0.05. See text or figure legends for more details.

## Author contributions

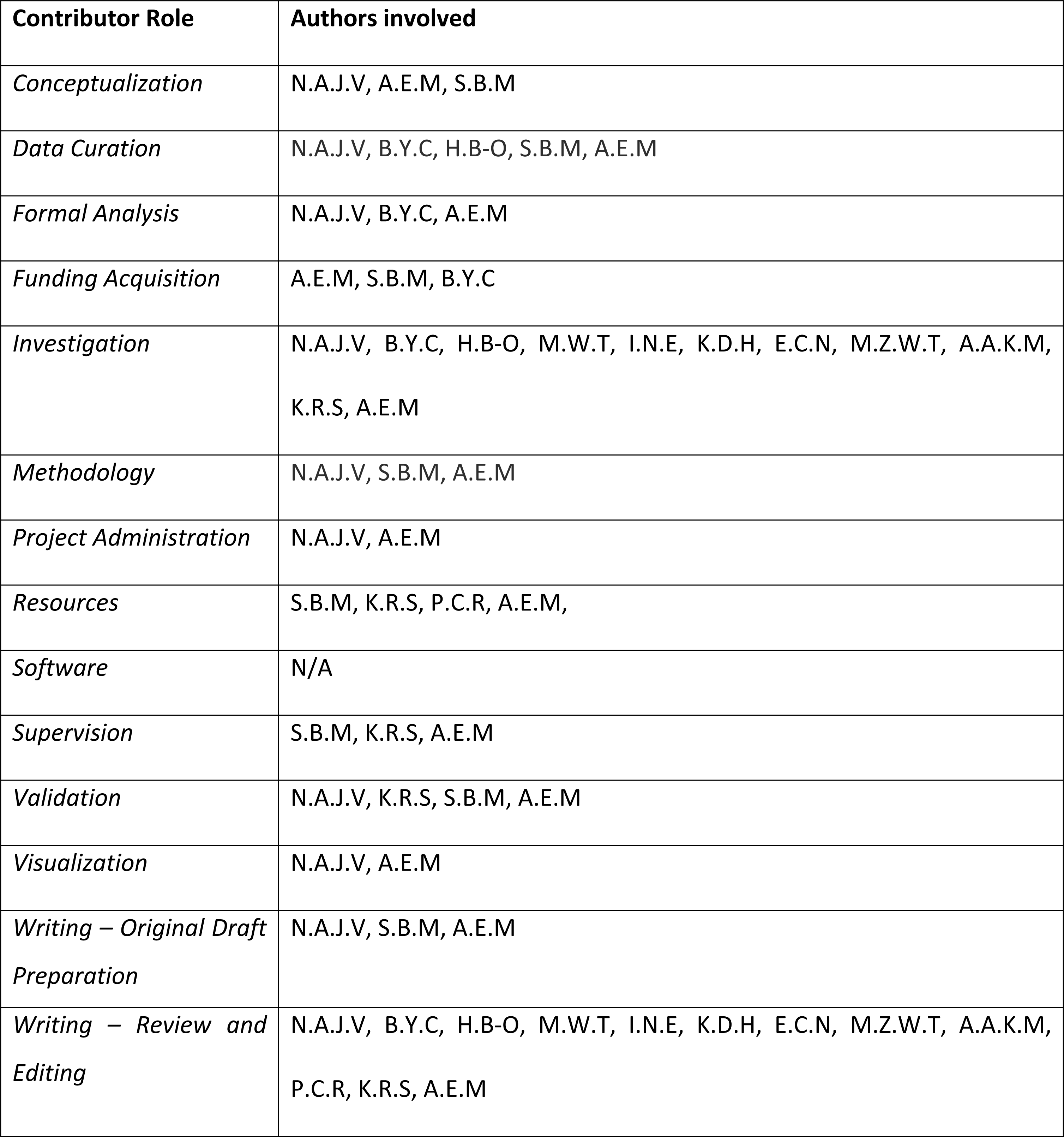

## Acknowledgements

This work was funded by grants to A.E.M and S.B.M (2020/GNT2002765) from the National Health and Medical Research Council of Australia and B.Y.C and A.E.M from the University of Melbourne, Australia. The pHW2000 plasmid used for quantifying IAV infection in tissue samples was sourced and distributed kindly from St. Jude Children’s Research Hospital. Cartoons were created with BioRender.com.

## Supplementary Figures

**Figure S1. *Body weight and lung cytokine measurements in vagotomized and sham mice following mock infection.*** Graphs depict body weight change and lung cytokine measurements in (A, C) right and (B, D) left vagotomized/ sham mice following intranasal mock (PBS) inoculation. N = 5 for each group at 8 days post infection. Data represented as mean ± SEM.

**Figure S2. *Body weight, clinical score, lung pathology and cytokine measurements in QX-314 and vehicle treated mice following mock infection.*** Graphs depict (A) body weight change, (B) clinic scoring, (C) lung histopathology and (D) lung cytokine measurements in mice receiving nebulized 300 μM QX-314 or vehicle (saline) following intranasal mock (PBS) inoculation. N = 10 for each group at days 4, 6, 8 post infection. Data represented as mean ± SEM.

**Figure S3. *Immunohistochemical analysis of MHC II expressing immune cells and CGRP expressing neurons within the vagal sensory ganglia of QX-314 and vehicle treated mice following mock infection.*** Graphs depict the percentage of (A) MHC II-expressing cells and (B) CGRP-expressing neurons per the total number of neurons in the vagal sensory ganglia in mice receiving nebulized 300 μM QX-314 or vehicle (saline) following intranasal mock (PBS) inoculation. N = 5 for each group at days 4, 6, 8 post infection. Data represented as mean ± SEM.

**Figure S4. Comparison of IAV H1N1 viral strains.** Graphs depict (A) lung cytokines, (B) lung viral titers, and (C) lung histopathological score post infection with either Auck/ 09 or PR8 H1N1 viral strains. N = 5-7 for each group at 3, 6, 8 days post infection. Data represented as mean ± SEM.

**Figure S5. *Flow cytometry gating strategy.*** Representative flow cytometry plots used to identify immune cell populations in mouse bronchoalveolar lavage fluid (BALF).

